# Dimerization of human PARP15 is required for NAD^+^ binding and automodification

**DOI:** 10.64898/2025.12.15.694324

**Authors:** Anna Tuovinen, Johan Pääkkönen, Mirko M. Maksimainen, Lea Hirschen, Heli I. Hentilä, Marie Tauscher, Bernhard Lüscher, Carlos Vela-Rodríguez, Patricia Korn, Lari Lehtiö

## Abstract

PARP proteins are enzymes catalyzing ADP-ribosylation, a well-conserved post-translational modification. In addition to the catalytic domain present in all PARPs, these proteins have a wide selection of other domains possessing different functions. Additional domains present in human PARP15 are macrodomains, capable of binding ADP-ribose. PARP15 is the least studied of the macrodomain-containing human PARPs, and it is linked to different diseases from infections to cancer. We show that the full-length canonical isoform 1 of PARP15 auto-ADP-ribosylates robustly on glutamate and aspartate residues, which can be hydrolyzed by known ADP-ribosylhydrolases. We have been able to locate several modification sites to the region of tandem macrodomains. In agreement with earlier reports regarding the catalytic domain dimerization being the requirement for enzyme activity, we show this in the full-length context with the recombinant isoform 1 of PARP15. We show that dimerization is required for the efficient substrate NAD^+^ binding, and we provide structural basis for this requirement with the help of a co-crystal structure of the catalytic domain dimer with an unhydrolyzable substrate analog.

## Introduction

ADP-ribosylation is an enzymatic covalent modification of proteins and nucleic acids, in which ADP-ribosyltransferases (ARTs) catalyze the transfer of ADP-ribose (ADPr) moieties from nicotinamide adenine dinucleotide (NAD^+^) to specific acceptors as nicotinamide gets released. The reaction results in mono-ADP-ribosylation (MARylation), involving the addition of a single ADPr unit, or in poly-ADP-ribosylation (PARylation), characterized by the formation of linear or branched ADPr chains. The human diphtheria toxin–like enzyme family of ARTs containing PARPs and tankyrases (TNKS1 and TNKS2) has 17 members, but only PARP1/2 and TNKS1/2 can catalyze PARylation while the others are MARylating enzymes or catalytically inactive proteins (PARP9/13) (Lüscher *et al*, 2022a; Vyas *et al*, 2014; Weixler *et al*, 2021). PARylation through PARP1 and PARP2 enzymes has been studied the most in the context of DNA damage repair. The PAR chains serve as scaffolds recruiting DNA repair proteins to the damaged site while also remodeling the chromatin through PARylation of histone tails to allow the repair to take place (Altmeyer *et al*, 2015; Leidecker *et al*, 2016; Messner & Hottiger, 2011). On the other hand, PAR chains can also act as a hindrance, inhibiting interaction physically or through repulsion (Zhu *et al*, 2025).

Studies of MARylation are significantly less encompassing compared to the long history of research on PARylation, but it is known that MARylation can act as an initiator of the PAR synthesis. One of the recognized roles of MARylation is to function as a signal in cells controlling protein interactions in various signaling pathways and in the interplay between pathogens and their hosts (Krieg *et al*, 2023; Lüscher *et al*, 2022b; Vyas *et al*, 2014). MARylation is recognized by macrodomains and ADP-ribosylhydrolases (ARHs) which are also able to remove it (Đukić *et al*, 2023; Forst *et al*, 2013; Jankevicius *et al*, 2013; Rosenthal *et al*, 2013). Recently, MARylation was reported as a site for ubiquitination resulting in a hybrid post-translation modification MAR ubiquitin ester (MARUbe) or mono-ADP-ribosyl ubiquitination (MARUbylation) (Bejan *et al*, 2024, 2025; Perrard *et al*, 2025; Pruneda *et al*, 2025).

In addition to a conserved catalytic domain, PARP enzymes consist of a selection of other domains, like WWE-domains, ARC-domains, KH-domains, zinc binding regions, and macrodomains thus diversifying the functions of these proteins (Suskiewicz *et al*, 2023). Macrodomains are evolutionarily conserved protein domains that are found in a wide range of organisms, having multiple recognized roles in cells from regulating gene expression at transcriptional level to chromatin remodeling and DNA repair (Aguiar *et al*, 2005; Costanzi & Pehrson, 1998; Han *et al*, 2011; Rack *et al*, 2016). They can bind ADP-ribosylated residues and function as readers or erasers of the modification, having higher affinity to either MAR or PAR, and also being selective towards the amino acid residue to which MAR/PAR is attached to (Forst *et al*, 2013; Karras *et al*, 2005; Weixler *et al*, 2025).

The significance and regulation of the MAR cleavage in the cells remains still poorly defined. There are macrodomains also in viruses that are known to erase the modification (Alhammad *et al*, 2021; Eckei *et al*, 2017; Li *et al*, 2016; McPherson *et al*, 2017). Hydrolysis by viral macrodomains has been seen to be targeted more efficiently towards MARylation rather than PARylation, highlighting the role of MARylating PARPs in innate immunity (Eckei *et al*, 2017; Lüscher *et al*, 2022b). For example, the viral non-structural protein 3 (nsp3) macrodomain from Chikungunya virus (CHIKV) can erase the MARylation from PARP10-mediated MARylation of the CHIKV protease (Krieg *et al*, 2023). The overexpression of macrodomains and hydrolase activity has also been observed in different cancer types and could thus be used as biomarkers in cancers diagnostics (Feijs *et al*, 2020; Han *et al*, 2011; Jankevicius *et al*, 2013; Palazzo *et al*, 2019).

PARP enzymes PARP9, PARP14 and PARP15 are equipped with macrodomains and are thus called macro-PARPs. Before identification as PARP proteins, PARP9 was found as a risk gene for diffuse large B-cell lymphoma and thus named BAL1 (B-aggressive lymphoma1) and based on the sequence similarity PARP14 and PARP15 were linked to the same protein family as BAL2 and BAL3, respectively (Aguiar *et al*, 2000). Macro-PARPs have been evolving under strong positive selection in primates and are therefore suspected to have an important role in viral defense and innate immunity. Gene loss and gain in PARP14 and PARP15 imply co-evolution with viruses, gene occurrence reflecting the need for immune defense at the time (Daugherty *et al*, 2014; Delgado-Rodriguez *et al*, 2023). Furthermore, PARP9 and PARP14 have been connected to macrophage activation, and their genes have been discovered to perform anti-inflammatory roles (Higashi *et al*, 2019; Iwata *et al*, 2016; Zhu *et al*, 2024).

Not much is known about the function of PARP15, but it has been suggested to be connected to cellular stress response, as it has been reported to localize to cytoplasmic stress granules (SGs) (Chan *et al*, 2024; Leung *et al*, 2011). SGs establish upon cellular stress, triggered for example by viral infections. In addition to PARP15, also TNKS1, PARP12, PARP13 and PARP14 are localized to SGs, from which at least PARP14 can work as initiator of PAR chains (Challa *et al*, 2025). The initial MARylation is further elongated to form PAR chains, which in turn supports the assembly of additional proteins to coordinate specific functions (Jayabalan *et al*, 2025; Lüscher *et al*, 2022b). PARP15 is also capable of ADP-ribosylating the phosphorylated ends of ssRNA, but whether this is targeted towards host or viral RNA is not known. Despite the common origin in evolution, PARP14 does not ADP-ribosylate RNA, suggesting unique roles for PARP15 instead of functioning as a backup for PARP14 (Munnur *et al*, 2019; Weixler *et al*, 2022). Recently, PARP15 was identified as a susceptibility locus for Clarkson disease, defined by leaking capillaries. In this study, the first physical target of PARP15 was found, as it was observed to ADP-ribosylate the scaffold protein JIP3 in human microvascular endothelial cells which inhibits the function of p38 MAP kinase pathway (Chan *et al*, 2024).

In contrast to other macro-PARPs, PARP15 is absent in several mammalian species, for example mice, which hampers *in vivo* studies of this protein (Shaw *et al*, 2017). It has two main isoforms that differ in their N-terminal regions; in isoform 1, which is the longer isoform, both macrodomains are present, while isoform 2 lacks the first macrodomain. In addition, isoform 2 has a unique N-terminus consisting of 20 amino acids. Some vertebrates express only isoform 2, but in humans, both isoforms are expressed (Daugherty *et al*, 2014). We have successfully purified the long isoform 1 of full-length (FL) PARP15 that will be referred to as “PARP15^FL^” from now on in this article. We show using activity and MAR detection assays, the latter employing the MAR binding macrodomain eAf1521 that PARP15^FL^ is an active enzyme that robustly MARylates itself. Moreover, we document, using site-specific hydrolases and mass spectrometry (MS), the location of several MARylated residues. We also observed that dimerization is required for efficient binding of the substrate NAD^+^ and for catalytic activity. We report a co-crystal structure of PARP15 with an unhydrolyzable substrate analog shedding light into the observed requirement of dimerization for enzymatic activity.

## Results

### Automodification and NADase activity of PARP15

We used the NAD^+^ hydrolysis assay to test the activity of PARP15^FL^ and the isolated catalytic domain of PARP15 (PARP15^CAT^). In this experiment, proteins are incubated with β-NAD^+^, and the remaining NAD^+^ is converted to a fluorophore that can be measured using 372 nm as excitation wavelength and 444 nm for emission (Putt & Hergenrother, 2004; Venkannagari *et al*, 2013). Both PARP15^CAT^ and PARP15^FL^ are active, and the consumption of NAD^+^ is boosted as the protein concentration is increased **(Fig. 1A)**. By using the macrodomain super binder eAf1521 fused to nanoluciferase we were able to visualize the intensity of the MARylation signal on western blot and determine if the consumption of NAD^+^ was targeted towards automodification (**Fig. 1B**) (Nowak *et al*, 2020; Sowa *et al*, 2021). On the membrane, a weak MARylation signal for PARP15^FL^ reveals that it is already slightly modified before the NAD^+^ incubation, originating from protein expression in insect cells Sf21. A robust increase of MARylation can be seen in PARP15^FL^ upon NAD^+^ incubation already at 1 µM protein concentration, whereas PARP15^CAT^ does not show an increase of MARylation in a similar fashion even when using 4 µM protein concentration, proposing that PARP15^CAT^ would act as NADase rather than ADP-ribosyl transferase in this assay. We also used MS to detect the increase of the molecular weight of PARP15^FL^ and PARP15^CAT^ after NAD^+^ incubation, corresponding to molecular weights of a single or multiple MARylations **(Fig. 1C–F)**. Here, PARP15^FL^ modifies itself as expected with a mass distribution ranging from non-modified to up to five modifications in a protein **(Fig. 1E)**. PARP15^CAT^ was also detected to modify itself (**Fig. 1F**), but the proportion of non-modified protein was high compared to the non-modified protein for PARP15^FL^, probably being the reason why the modification signal was so low on the western blot.

**Fig 1.**
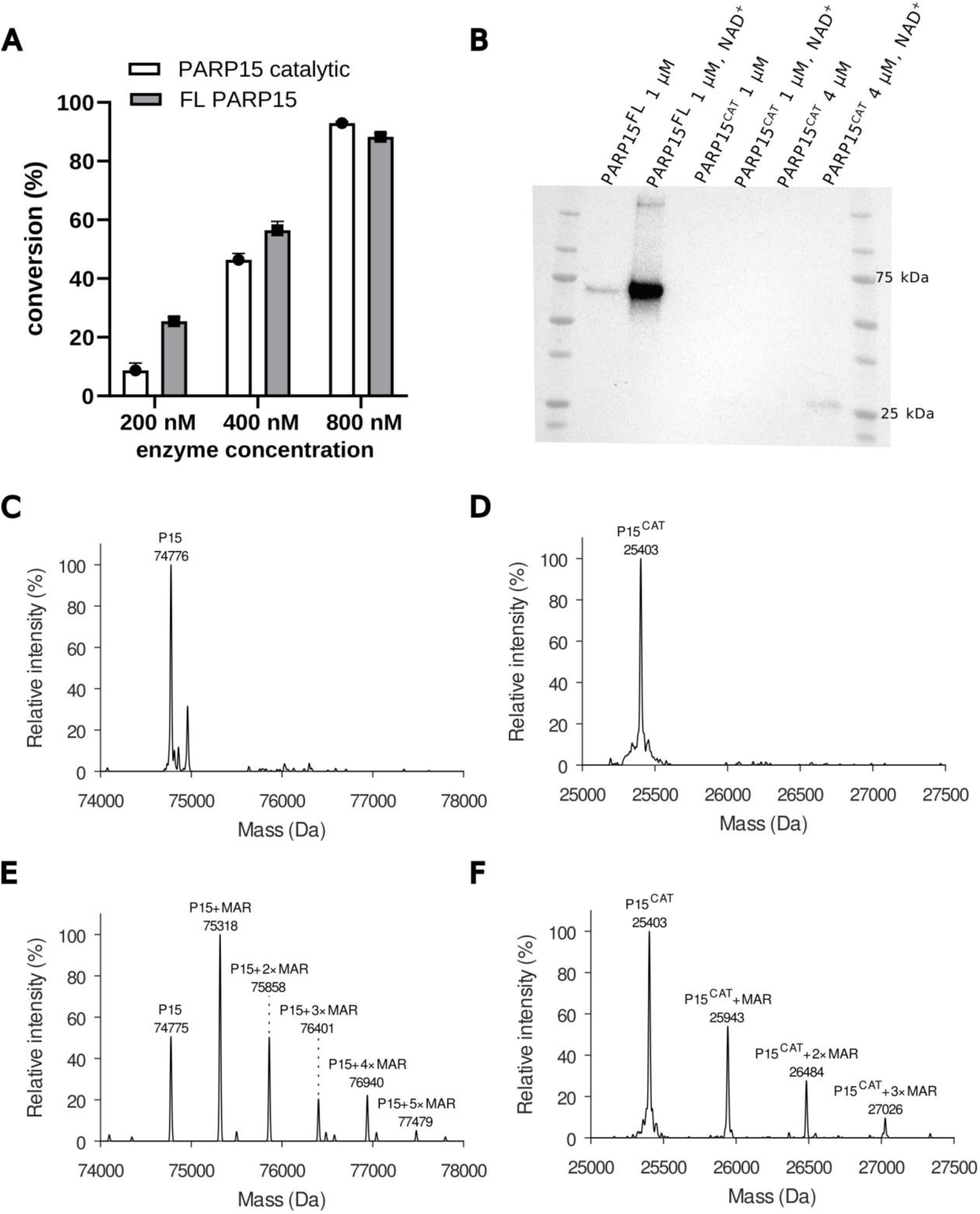
Enzyme activity and automodification of PARP15^FL^ and PARP15^CAT^. A) NAD^+^ conversion assay for PARP15^CAT^ and PARP15^FL^ showing that both enzymes are active hydrolases. B) In western blot PARP15^FL^ automodification is strongly increased upon NAD^+^ hydrolysis, whereas PARP15^CAT^ MARylation band is weak. C) Deconvoluted mass spectrum of PARP15^FL^ in denatured state shows that the protein is all unmodified. D) Similarly, PARP15^CAT^ is unmodified after purification. E) After NAD^+^ incubation, PARP15^FL^ automodifies to form a mass distribution with 0–5 detectable MARylations. F) Similarly, PARP15^CAT^ automodifies and 0–3 MARylations are detected.

### PARP15 ADP-ribosylates itself on acidic residues

As there was an increase in the PARP15^FL^ automodification after NAD^+^ incubation, a panel of macrodomains and ADP-ribosyl hydrolases with known target specifity was tested to uncover to which amino acid residues MARylation is attached (**Fig. 2A**). Human MacroD1 (MDO1) and MacroD2 (MDO2) are capable of cleaving the established ester bond when MAR is attached to aspartate or glutamate (Agnew *et al*, 2018; Neuvonen & Ahola, 2009; Rosenthal *et al*, 2013). Recently, also the first macrodomains in PARP9 and PARP14 have been reported to possess hydrolysis activity towards MARylated acidic residues (Đukić *et al*, 2023; Weixler *et al*, 2025). Similarly, the SARS-CoV-2 nsp3 macrodomain represents a viral macrodomain that can hydrolyze MAR attached to Asp/Glu (Alhammad *et al*, 2021). Besides acidic residues being MARylated, modification can be attached to arginines, which is reversed by the action of ARH1, or to serines, which is recognized and hydrolyzed by ARH3 (Abplanalp *et al*, 2017; Fontana *et al*, 2017; Moss *et al*, 1985; Oka *et al*, 2006). MacroH2A.1.1, by contrast, is not expected to be active in hydrolyzing MARylation (Forst *et al*, 2013; Jankevicius *et al*, 2013). We also tested PARP15 macrodomain 2, which we assumed would be rather a reader than eraser of MARylation based on the close evolutionary origin and similarity to PARP14 macrodomain 3, recognized as a reader of MARylation (Forst *et al*, 2013; Haikarainen *et al*, 2018). If the MARylation could not be cleaved using these site-specific hydrolases, it would point out that MARylation occurs on some of the previously identified possible MARylation sites like cysteine, lysine or histidine, for which hydrolases are still missing (Minnee *et al*, 2024).

**Fig 2.**
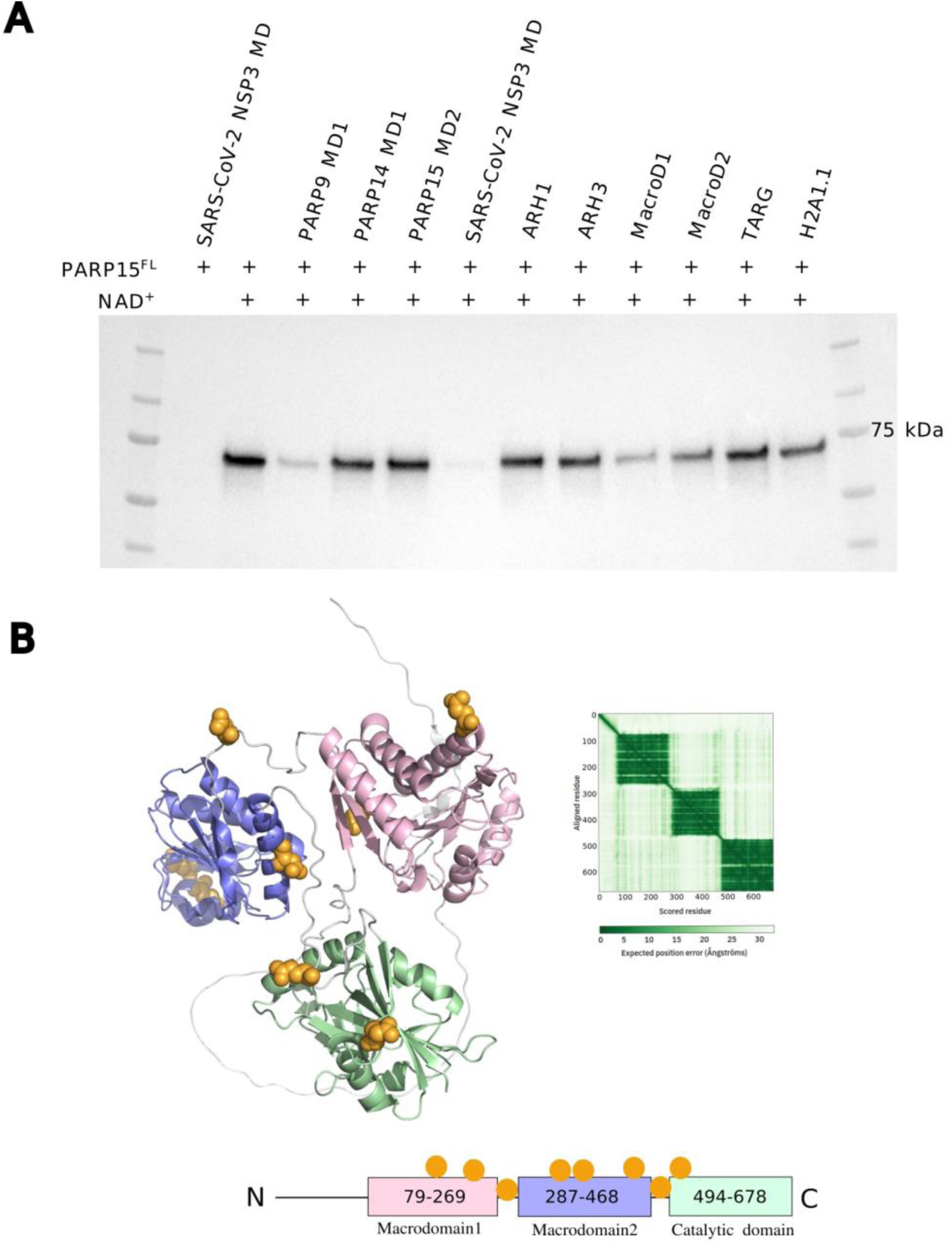
PARP15^FL^ automodifications in acidic residues can be removed with site-specific macrodomains. A) MARylation of PARP15^FL^ expressed in and purified from insect cells can be cleaved with SARS-CoV-2 nsp3 macrodomain, suggesting that only Asp/Glu are modified in this cell system. PARP15^FL^ modifies itself upon NAD^+^ incubation and PARP9 macrodomain 1, SARS-CoV-2 nsp3 macrodomain and human macrodomain 1, which are site-specific to aspartate and glutamate, are able to erase the MARylation. B) AlphaFold prediction of the PARP15^FL^ suggests that the protein consists of a long unstructured N-terminal tail followed by three domains (two macrodomains and the catalytic domain) having short flexible regions between them. There is no predicted interaction between the domains. Identified MARylation sites are located in the domains and also between them in the flexible regions, marked as spheres.

To evaluate this panel of hydrolases, we first allowed automodification of PARP15^FL^ by incubating the enzyme with NAD^+^ followed by hydrolysis of MARylation with a macrodomain or hydrolase. As mentioned earlier, PARP15^FL^ is MARylated during expression in insect cells, and we discovered that this modification could be completely removed by pretreating it with SARS-CoV-2 nsp3 macrodomain. This indicates that likely only Asp and Glu are modified during protein expression. After NAD^+^ incubation, a clear decrease of MARylation was observed with some macrodomains that are targeted towards MAR cleavage from acidic residues, revealing that the most prominent site of the modification is Asp and/or Glu (**Fig. 2A**), following the same trend already reported for PARP15^CAT^ (Wallace *et al*, 2021). PARP14 macrodomain 1 was not seemingly cleaving MARylation even when using twice as high concentration (4 µM) as for the other cleaving macrodomains (2 µM). It is, however, not reported if the sequence of target proteins around MAR sites is important to specify the sensitivity to hydrolases, i.e. co-determining binding specifity and thus catalytic activity. This could be due to steric hindrance inhibiting the hydrolytic activity for certain enzymes, pointing out this as a possible explanation for the divergent hydrolytic activity for the enzymes supposed to target the same modified residues. TARG1 is also reported to be site-specific for acidic residues, but we did not observe a decrease in the MARylation signal with PARP15^FL^, which on the other hand, is in line with our previous observations (Haikarainen *et al*, 2018; Sharifi *et al*, 2013).

As we identified biochemically that self-MARylation in PARP15^FL^ mainly targets acidic residues, we focused on mapping those sites using bottom-up proteomics. After pretreatment with SARS-CoV-2 nsp3 macrodomain to remove the MARylation that occurs in insect cells, subsequently we incubated PARP15^FL^ with NAD^+^ and stabilized the modifications using hydroxylamine which attacks the ester group and leaves a stable hydroxamic acid group on the modified residue (15 Da larger than unmodified carboxylic acid) (Gomez *et al*, 2018; Wallace *et al*, 2021; Zhang *et al*, 2013). Using this approach, we identified several automodification sites in PARP15^FL^; hydroxamic acid groups were found in E177, E243, D281, E348, E350, D458, E484 and E500 (**Fig. 2B**) in three independent analyses. All these sites were of low occupancy, proposing that in PARP15^FL^ there are multiple modification sites that end up only partially MARylated, supported by the results from intact mass distribution of PARP15^FL^ **(Fig. 1E).**

### Dimer of PARP15 is the catalytically active form of PARP15^FL^

Previously, we have already seen that PARP15^CAT^ is a dimer in solution in contrast to PARP10^CAT^ being a monomer (Venkannagari *et al*, 2013). Recently, guided by multiple crystal structures available for PARP15^CAT^, residues R576 and D665 in the dimerization interface were mutated resulting in a decrease in the enzymatic activity (Ebenwaldner *et al*, 2025). We confirmed these results using NAD^+^ hydrolysis assay for PARP15^CAT^ D665A, D665R and R576A showing that these mutants were clearly not as active as wild type (WT) PARP15^CAT^ **(Fig. 3A)**. We also confirmed with native MS that the CAT domain forms a dimer whereas the mutations at the interface prevent dimerization (**Fig. S1**). We then produced mutants D665A and D665R also in PARP15^FL^ and we noticed that the mutation drastically decreases enzyme activity also in FL protein **(Fig. 3A)**. The automodification ability of these enzymes was also shown to be halted on western blot, showing only MARylation signal originating from expression in insect cells Sf21, but not showing an increase upon NAD^+^ incubation **(Fig. 3B).** The dimer formation indeed seems to be the activation mechanism for this enzyme also in the FL context; in size-exclusion chromatography with multi-angle light scattering (SEC-MALS) PARP15^FL^ D665A and D665R clearly elute as monomers whereas molecular weight (MW) for PARP15^FL^ WT is closer to the size of a dimer **(Fig. 3C)**. Masses calculated at the maxima of the chromatogram peaks for D665A was 71.53 kDa and 74.00 kDa for D665R, corresponding to theoretical mass 74.73-74.82 kDa quite well. For the dimer, despite clearly forming a higher oligomer, the measured MW is not stable over the peak resulting in a lower average MW (122.6 kDa) than expected. The unstable MW estimate is also visible in the asymmetric form of the elution peak.

**Fig 3.**
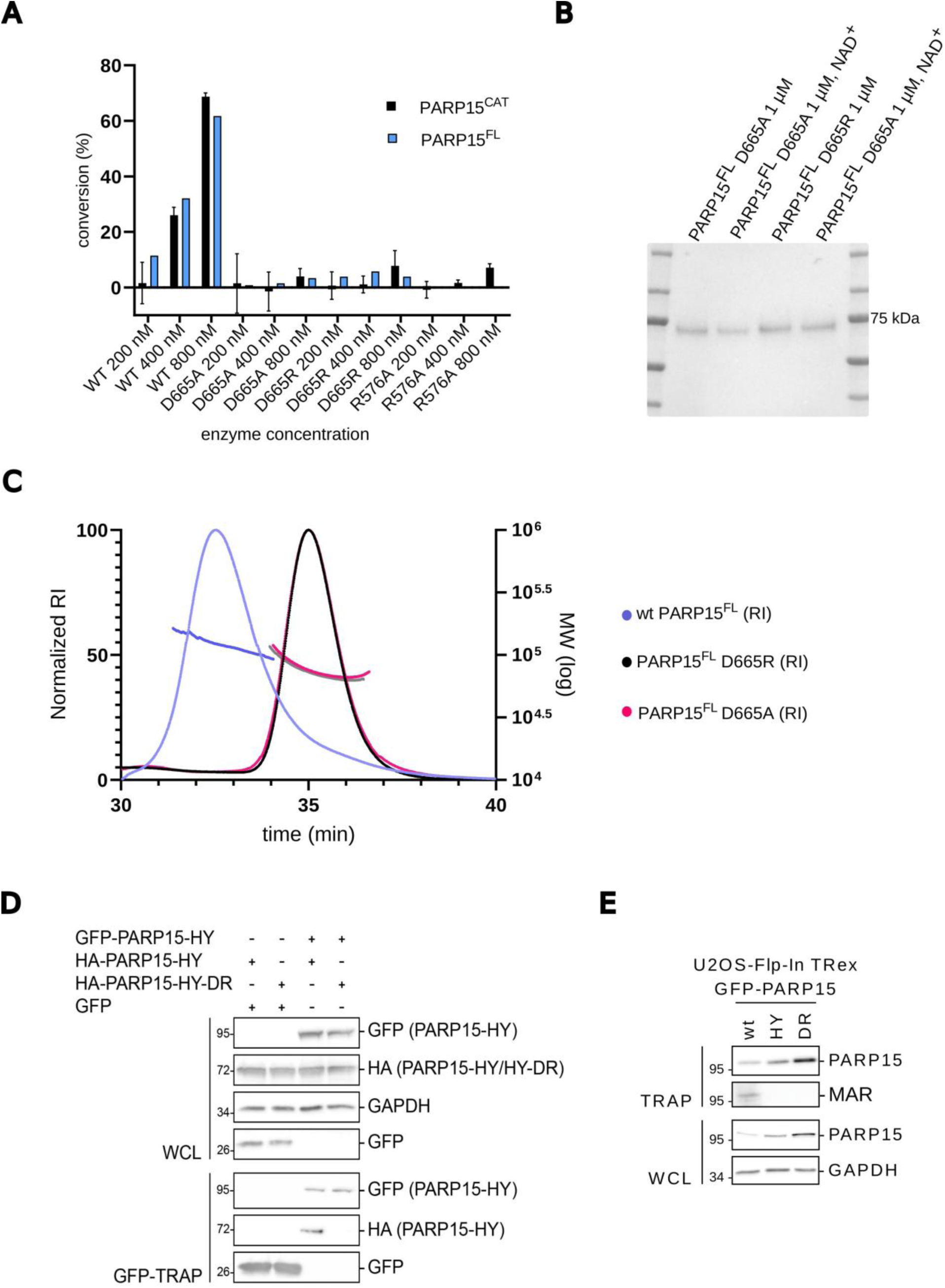
Dimer formation is a requirement for active PARP15^FL^. A) A difference in activity is seen between the WT and mutant proteins using NAD^+^ hydrolysis assay, showing that both PARP15^CAT^ and PARP15^FL^ D665A, D665R and PARP15^CAT^ R576A mutants are not hydrolyzing NAD^+^ efficiently. B) PARP15^FL^ D665A and D665R are modified during expression in the Sf21 insect cells, showing MARylation before the incubation with NAD^+^ *in vitro*, but not showing enhanced MARylation upon NAD^+^ incubation. Full western blot available in Supplementary **(Fig. S2)**. C) In SEC–MALS PARP15^FL^ WT elutes as a dimer, whereas D665A and D665R elute as monomers. D) PARP15^FL^ H559Y tagged with either GFP or HA and HA-PARP15^FL^ H559Y D665R were expressed in HEK293 cells. GFP-tagged proteins were enriched using GFP-TRAP. Co-purified HA-tagged proteins were evaluated on western blots. For control, whole cell lysates (WCL) were analyzed for expression of the indicated proteins. E) The expression of GFP-PARP15^FL^, GFP-PARP15^FL^ H559Y and GFP-PARP15^FL^ D665R was induced with doxycycline in U2OS cells and subsequently the indicated proteins were enriched using GFP-TRAPs. PARP15 expression was monitored using specific antibodies after enrichment and in WCL. MARylation was assessed in the enriched samples.

In the PARP15 catalytic cleft, conserved amino acid triad HYL is needed for the enzyme activity (Challa *et al*, 2021). For the cell co-immunoprecipitation experiments, we mutated one of these amino acids and produced an inactive PARP15^FL^ enzyme H559Y (PARP15-HY) to monitor the dimer formation independently from enzymatic activity (Karlberg *et al*, 2015; Korn *et al*, 2025). We used distinctly tagged versions of PARP15^FL^ to test interaction in cells. Using GFP-trap, GFP-tagged PARP15^FL^ HY co-precipitated HA-tagged PARP15^FL^ HY but not the dimerization mutant PARP15^FL^ D665R. This indicates that the binding interface is the same *in vivo* as *in vitro,* and the role for residue D665 is significant for the formation of oligomer **(Fig 3D)**. We also show that PARP15^FL^ is not able to catalyze ADP-ribosylation as a monomer in the cells; PARP15^FL^ D665R was not MARylating, and the activity of the protein was comparable to the inactive PARP15^FL^ HY-mutant **(Fig 3E)**. However, evaluating the modification activity in the cells is not straightforward, because there is no existing data about the magnitude of automodification of PARP15 in cells. Based on our findings, we propose that PARP15^FL^ is not a monomer in the cells, and the ability to form an oligomer is not dependent on the catalytic activity of the protein, but on the other hand, the oligomer formation appears to be a prerequisite for the active enzyme.

### Inter- and intradomain interactions within the PARP15 dimer

The AlphaFold model indicates that for PARP15^FL^ the CAT domain is the main contributor for the dimer formation and there is very little support for a contribution of the macrodomains and the N-terminal unstructured region, the prediction of which is of very low pLDDT (predicted local distance difference test) (**Fig. 4A**). Also for isoform 2, CAT domains are predicted to form the only dimer interface **(Fig. S3)**. The modification sites we mapped to the tandem macrodomain region are in line with the dimer model as the glutamate and aspartate residues are accessible on the molecular surface **(Fig. S4)**. In isoform 1, macrodomains are predicted to form a single unit having interactions between the macrodomains of the same PARP15 monomer. The prediction is in line with our observation that D665R point mutation would be enough to dissociate the dimer in PARP15^FL^. To further study the AlphaFold-generated dimer model, we carried out hydrogen-deuterium exchange mass spectrometry (HDX-MS). In contrast to the AlphaFold model, we observed that there is a clear change in the deuteration rates also in macrodomains when comparing the WT and the dimerization mutants D665A and D665R. The macrodomains and the catalytic domain in the PARP15^FL^ mutants are more exposed to solvent as evidenced by higher deuteration rates and suggesting partial unfolding, whereas the domains are less flexible in WT as the conformation is more packed **(Fig 4B)**. The deuteration rates were determined at four time points, the first of which at 20 seconds had the clearest differences between WT and mutants **(Fig. S5)**. In contrast, the deuteration rates converged in most regions by 2 hours, presumably because WT was slowly destabilized as the dimer dissociated in the dilute deuteration condition. This indicates that besides the region in the catalytic domain close to the dimerization interface, several regions in the macrodomains are affected by the interactions in the dimer, and dimerization overall leads to a more compact structure of the protein.

**Fig 4.**
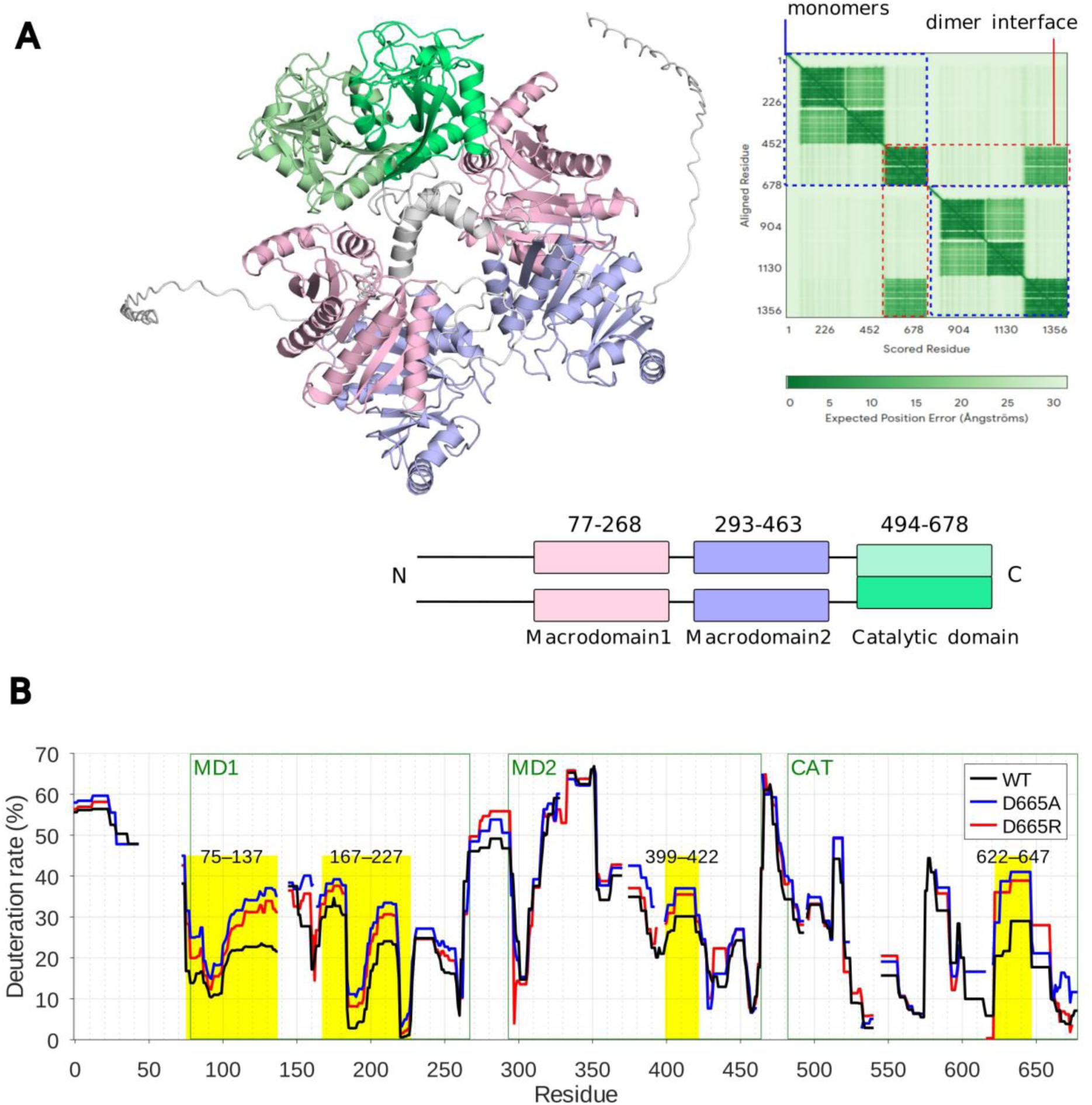
PARP15 dimer formation changes the domain dynamics. A) AlphaFold dimer model predicts that in PARP15^FL^ the interface is only between the catalytic domains. B) Deuteration rates of PARP15^FL^ WT, D665A and D665R after 20-second deuteration time in HDX-MS show that several regions (highlighted in yellow) of D665A and D665R are deuterated more quickly and are therefore more flexible than the corresponding regions in WT dimer. Deuteration plots of representative peptides in the flexible regions are shown in **Fig. S5**.

### Dimerization is needed for substrate binding of PARP15

When we added the unhydrolyzable NAD^+^ analog benzamide adenine dinucleotide (BAD) to PARP15^CAT^ WT, D665A and D665R in 10:1 molar ratio, monomeric forms of the protein whether WT or the interface mutant did not bind BAD efficiently **(Fig. 5A)**. The dimeric form of WT bound BAD with 1:1 stoichiometry, and only minute amounts of apo-form of the dimer were detected **(Fig. 5B)**. We then quantified the affinity of BAD to the PARP15^CAT^ WT dimer by preparing and measuring titration series, which yielded the dissociation constant *K*_d_ = (29 ± 2) µM and maximum specific binding *B*_max_ = 0.99 ± 0.02 (mean ± standard deviation, *n* = 3) **(Fig. S6)**. The apparent amount of the PARP15^CAT^ monomer–BAD complex was so low that its *K*_d_ could not be reliably determined.

**Fig. 5.**
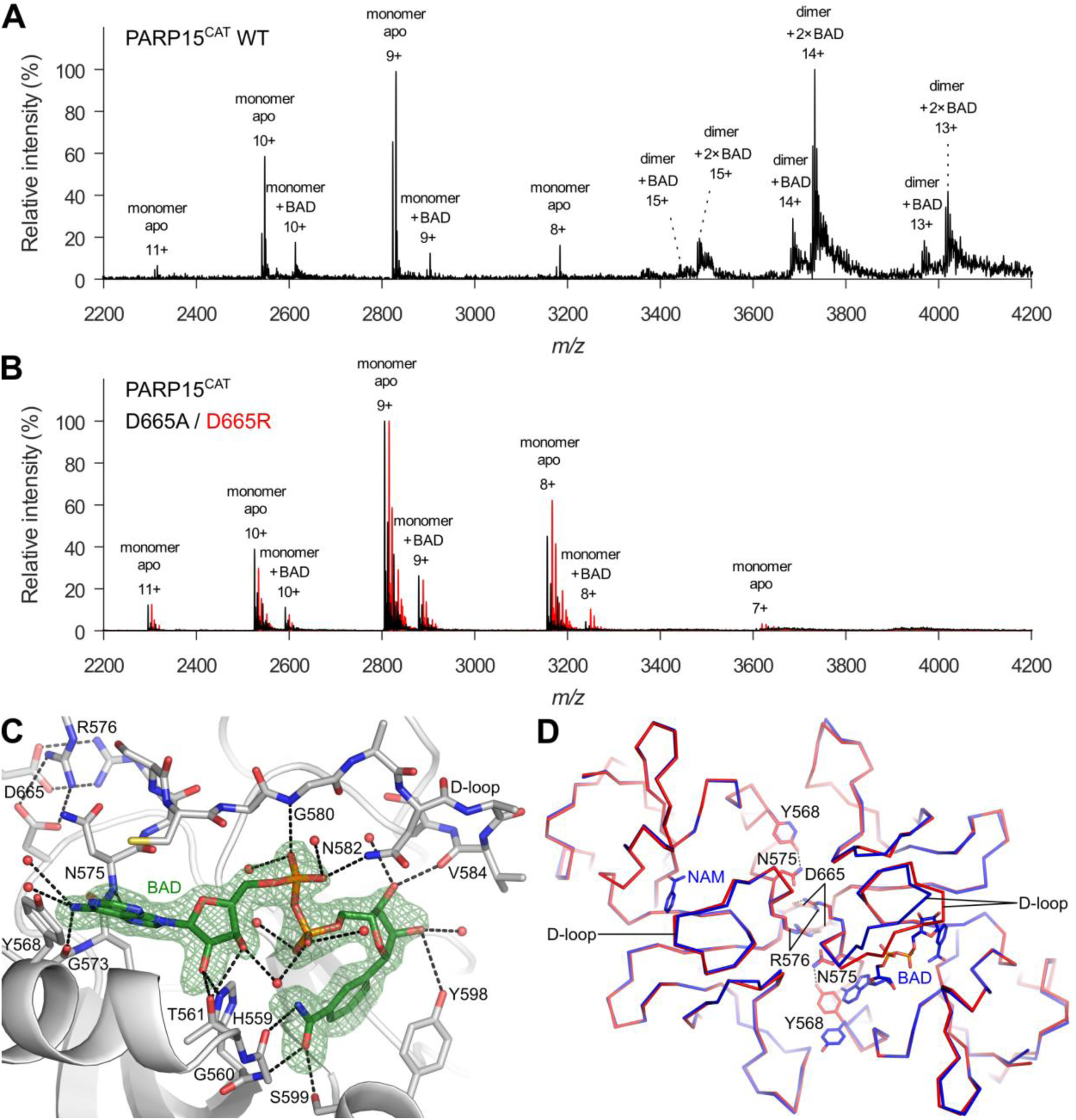
PARP15CAT binding to NAD^+^-analog BAD. A) The native mass spectrum of 10 µM PARP15^CAT^ WT + 100 µM BAD shows that the PARP15^CAT^ dimer binds BAD much more strongly than the monomer does. B) Native mass spectra of 10 µM PARP15^CAT^ D665A (black) and D665R (red) + 100 µM BAD show that the mutants behave like the WT monomer, i.e. they bind BAD only weakly. C) BAD bound to the catalytic site in the PARP15^CAT^-BAD crystal structure. Hydrogen bonds to the protein and water molecules are shown as dashed lines. The homodimer is formed via two inter-chain salt bridges between R576 and D665, also shown in the top-left corner. The *mF*_o_−*DF*_c_ omit map of BAD is shown with a 3*σ* contour level. **D)** The PARP15^CAT^-BAD homodimer (blue) superimposed with the PARP15^CAT^ apo-form (chain B of pdb_00003blj, red) shows how the binding of BAD in chain A (right) changes the conformations of the D-loop and the side chain of Y568 in comparison to chain B (left) where only nicotinamide (NAM) is bound. In this flipped conformation, Y568 does not form a hydrogen bond with N575.

We solved the crystal structure of PARP15^CAT^ with BAD **(Fig. 5C**, **Table 1**) and compared it to the apo-structure (pdb_00003blj)(Karlberg *et al*, 2015) showing a change in the D-loop upon binding of BAD (**Fig. 5D**). Comparison of our dimer PARP15^CAT^ structure between the A and B chain reveals the conformational change upon BAD binding; when BAD is accommodated in the binding pocket of one chain, the D-loop in the other binding pocket is in a different conformation. Also, Tyr568 is pointing to the almost completely opposite direction inside the binding pocket partially filling the pocket. Based on the orientation and 3.2 Å distance to Asn575 there might be a hydrogen bond stabilizing the tyrosine orientation in the closed conformation. As the binding affinity of the monomer to BAD is low compared to the WT dimer, our hypothesis is that there are certain interactions in the dimer to control the opening of the D-loop to accommodate BAD that are missing in the monomer.

**Table 1.**
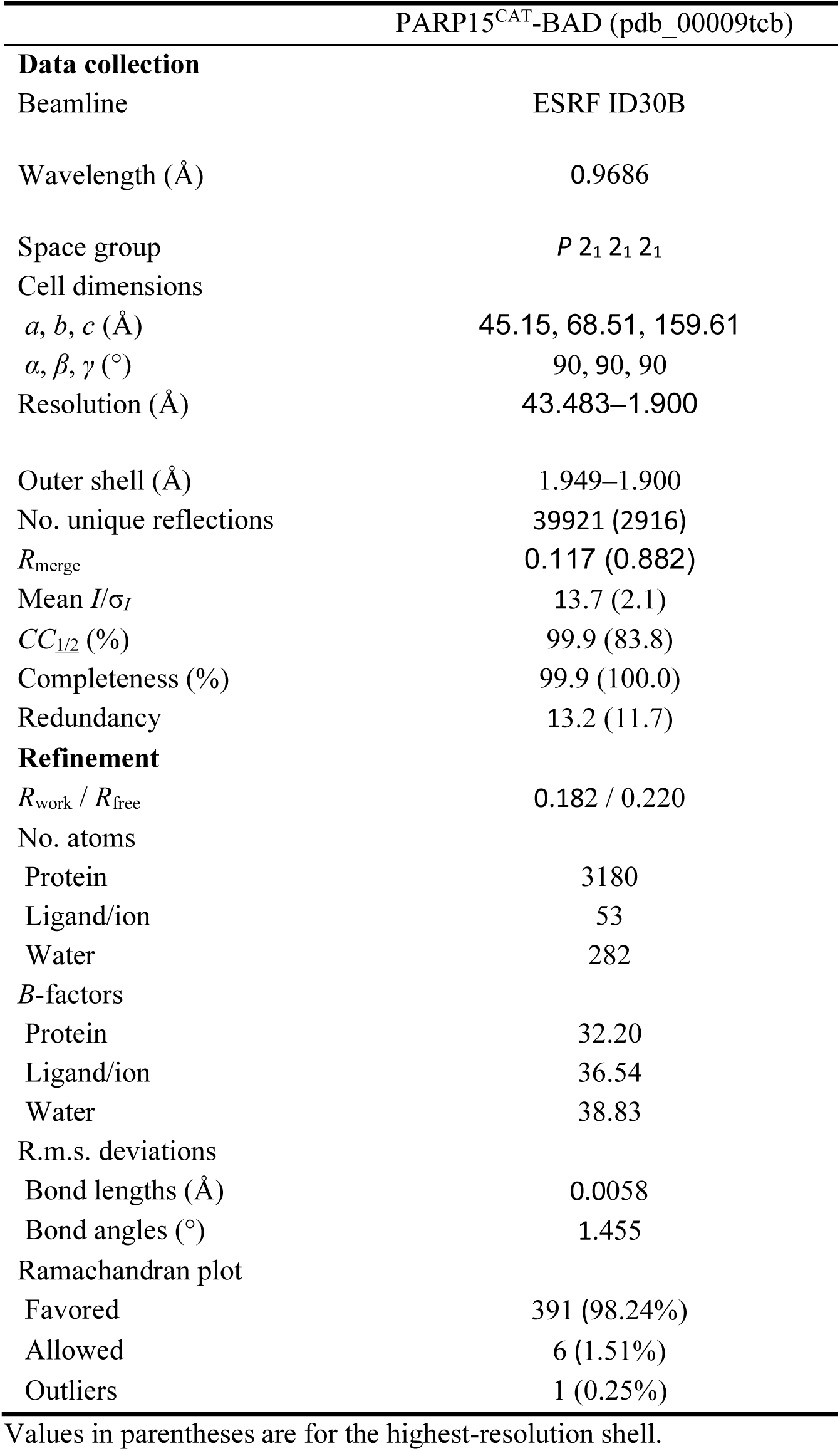
Data collection and refinement statistics.

## Discussion

Enzymes often exist as higher order oligomers, and oligomerization has been identified as a key factor in their regulation (Marianayagam *et al*, 2004). Using recombinantly expressed proteins, we show that dimer formation is critical for the activity of human PARP15 both for the isolated catalytic ART domain (PARP15^CAT^) as well as for the full-length long isoform (PARP15^FL^). While both constructs showed robust consumption of NAD^+^ **(Fig. 1A)**, PARP15^CAT^ appeared to mainly function as an NADase with little automodification activity while PARP15^FL^ strongly automodified **(Fig. 1B)**. In agreement with this, we demonstrate using mass spectrometry that while we can detect PARP15^CAT^ automodification, in the context of PARP15^FL^ the automodification frequency is substantially higher (**Fig. 1E and F**). There is a need for a regulatory mechanism of PARP15^FL^ in cells, suggested by the fact that NAD^+^ cannot be consumed uncontrollably as it would be harmful for the cells because NAD^+^ is important for many key processes, such as glycolysis (Hopp *et al*, 2019). While we can detect automodification of PARP15 in cells, this only corresponds to the automodification activity of the homodimer (**Fig. 3E**). Other ways to regulate PARP15 activity would be through target protein proximity and engagement and by the availability of NAD^+^ that can be different between different cell compartments (Ryu *et al*, 2018).

MARylation is important for innate immune defense, and macrodomains encoded by viruses can counteract this. Virus macrodomains hydrolyze MARylation and thereby control the protein composition in SGs as part of their mechanism to evade the host immune response (Bonenfant *et al*, 2019; Jayabalan *et al*, 2021, 2025; Peng *et al*, 2008). We mapped the main auto-MARylation sites in PARP15^FL^ mainly to the tandem macrodomains (**Fig. 2B**) and also observed that SARS-CoV-2 nsp3 macrodomain efficiently hydrolyzes PARP15 MARylation consistent with the role of PARP15 in innate immunity. As the MARylation occurs in acidic residues, PARP9 macrodomain 1 was also seen to hydrolyze the modification, which is in line with the previous reports about PARP9 macrodomain 1 residue preference for MAR hydrolysis (Đukić *et al*, 2023; Weixler *et al*, 2025)(**Fig. 2A**). Also previously, acidic residues have been found as the main modification sites *in vitro* and *in vivo* among PARP enzymes (Buch-Larsen *et al*, 2025; Daniels *et al*, 2014; Gomez *et al*, 2018; Wallace *et al*, 2021; Zhang *et al*, 2013). The hydrolysis of MARylation remains largely unexplored, especially regarding the responsible enzymes. In human cells, there are several macrodomains that target similar residues, while they are also located in different compartments. As an example, MDO1 and MDO2, which also hydrolyze MARylations from acidic residues, are in mitochondria and in cytosol or nucleus, respectively (Agnew *et al*, 2018; Žaja *et al*, 2020). We did not observe self-hydrolysis of PARP15 MARylation by the PARP15 macrodomain 2, although it has been reported to be active and hydrolyze ADP-ribose from cysteine in pertussis toxin (Weixler *et al*, 2025). While we were able to map several modified residues in PARP15^FL^ *in vitro* using mass spectrometry, the role of these sites is not yet clear. The modifications could provide an additional regulatory layer for the enzyme, recruit in a MARylation-dependent manner target proteins to be further ADP-ribosylated or they could even act as initiators for PARylation in certain contexts like forming a platform for PAR-binding proteins in SGs.

While the dimer structure of PARP15^CAT^ has been observed in a wealth of crystal structures and observed for other PARPs (Jessop *et al*, 2024; Karlberg *et al*, 2015), our view on the dimerization of the PARP15^FL^ relies on the prediction (**Fig. 4A**). The PARP15^CAT^ lost the activity when the dimer interface was disrupted by site directed mutagenesis and this was also observed in the FL protein context, both in vitro and in cells (**Fig. 3**), suggesting that the dimer interface in isolated catalytic domains is also the dimer interface for PARP15^FL^. This is in agreement with recently reported findings for the short PARP15^FL^ isoform 2 (Ebenwaldner *et al*, 2025). The AlphaFold model also shows that there would be additional intra-protein contacts between the PARP15 macrodomains. With HDX-MS we were able to confirm this observation experimentally to some extent as there are several more flexible regions in monomeric mutants D665A and D665R in comparison to the dimerizing wild type enzyme (**Fig. 4B**). This could also have an effect on the MARylation reading function of the macrodomain and direct it towards specific so far unidentified targets.

In the apo form of the catalytic domain, the D-loop, while typically flexible in the crystal structures, as judged by weak electron density and high atomic displacement parameters, also partially closes the NAD^+^ binding site. In the monomeric form of the catalytic domain, the loose conformation could drive the changed D-loop dynamics towards a more collapsed conformation, and thus the protein would not be able to bind the substrate NAD^+^. Upon binding of the substrate analog BAD the loop opens and adopts a distinct conformation (**Fig. 5D**). The dynamics of the D-loop near the dimer interface is likely responsible for the substrate binding as a monomeric form lacking the supporting interactions in the beginning of the D-loop with the second subunit. There is a striking difference in the native mass spectra demonstrating that only the dimeric form of the PARP15^CAT^ can bind the BAD in a 1:1 stoichiometry confirming the results of the interface mutants (**Fig. 5A**). Here our interpretation differs from a recent report that dimerization would not affect substrate binding but target engagement (Ebenwaldner *et al*, 2025). Although the dimerization likely would affect binding to macromolecular targets that are to be ADP-ribosylated, the NAD^+^ consumption assay, where the PARP15^CAT^ acts mostly as an NADase, together with the observation that BAD binding occurs efficiently only to the dimeric form of the enzyme support our view that substrate NAD^+^ binding is impaired in the monomeric form of the enzyme. In the HDX-MS data, we observed that the loop region 622–647 in the CAT domain has higher deuteration rates in a monomeric form due to the contacts lost between the CAT domains (**Fig. 4B**). While dimerization is required for efficient substrate NAD^+^ binding, other structural changes induced by target protein binding could provide an additional level of interaction and context-dependent regulation of PARP15.

## Materials and methods

### Cloning

For the production of recombinant proteins, pNIC28-Bsa4 vector (#26103) was used for PARP15^CAT^ WT, R576A, D665A and D665R, and pFastBac-His-MBP (#30116) for PARP15^FL^ WT, D665A and D665R. DNA inserts with the correct mutation sites were first generated in PCR by having WT protein construct as a template and primers spanning over the mutation site as forward and reverse in separate reactions, and then combining the products to a single reaction with primers over the complete sequence, having appropriate overhangs for the vector. Vectors were linearized by amplifying the vector backbone, after which SLIC method was used to combine the linearized vector to insert, and the constructs were transformed into bacteria cells NEB5α or XL1 Blue, grown at 37°C on Luria Bertani (LB) agar plates supplemented with kanamycin (50 µg/ml) and sucrose (5% w/v) (Jeong *et al*, 2012). Small LB cultures supplemented with 50 µg/ml kanamycin and one inoculated colony were incubated overnight at 37°C and 220 rpm. DNA was extracted with Macherey-Nagel’s plasmid and DNA purification kit (NucleoSpin Plasmid EasyPure).

For constructs in the cells, HA- and GFP-PARP15 fusions were created using Gateway cloning. pDONR221-PARP15.1-HY vector was used for the transfection of an inactive PARP15 mutant (Korn *et al*, 2025). This was also used as a template for site directed mutagenesis.

The correct sequences were confirmed by sequencing. Details for cloning of the other DNA constructs used in this article are described in Supplementary Table 1.

### Protein expression

PARP15^CAT^ constructs were transformed to *E. coli* BL21(DE3) and grown on LB agar plates at 37°C supplemented with kanamycin (50 µg/ml) and sucrose (5% w/v). Small LB pre-cultures with inoculated colonies supplemented with kanamycin (50 µg/ml) were incubated overnight at 37°C in a shaking platform. The pre-cultures were used to inoculate autoclaved autoinduction media terrific broth base including trace elements (Formedium) with a 1:100 dilution. The expression cultures were supplemented with kanamycin (50 µg/ml) and glycerol (8 g/L) and incubation was continued until OD_600_ was 1. The temperature was shifted to 18 °C and grown O/N. The cultures were harvested by centrifugation and cell pellets were resuspended to lysis buffer supplemented with 0.02 mg/ml DNase and 0.1 mM Pefabloc® (Sigma-Aldrich). The biomass was flash-frozen with liquid nitrogen and stored at −20°C before the protein was purified.

Insect cell line Sf21 was used for the expression of FL protein constructs using Bac-to-Bac® baculovirus expression system (Invitrogen). Transfection of recombinant bacmids to Sf21 cells was performed using FuGENE®6 (Promega E2693) and V_0_ virus was collected after 7 days of incubation at 27°C. Virus was further amplified and collected after 5 days (V_1_). The large expression culture at 0.8 million cells/ml was infected with V_1_ and after 5 days harvested using centrifugation. Cells were resuspended to lysis buffer supplemented with Pierce protease inhibitor (Thermo Scientific A32963), and biomass was flash-frozen with liquid nitrogen and stored at −20°C before the protein was purified. Details for protein expression of the other proteins are described in Supplementary Table 1.

### Protein purification

Cells were lysed by sonicating using a Branson digital sonifier 450 followed by separation of soluble protein from cell debris by centrifugation. Supernatant was filtered through 0.45 µm filter for proteins expressed in *E. coli*, but for proteins expressed in Sf21 an additional centrifugation step was carried out instead. All the proteins contain His-tags and were therefore purified using immobilized metal affinity chromatography (IMAC). Prepacked IMAC column (Cytiva) loaded with Ni^2+^ was used for the proteins expressed in *E. coli* (PARP15^CAT^). HisPur™ Ni-NTA Resin (ThermoScientific) was used for proteins expressed in insect cells Sf21(PARP15^FL^) followed by further polishing with Amylose resin (New England Biolabs). *E. coli* proteins were purified at RT and insect cell proteins at +4°C. The columns were washed with lysis, wash (with an additional wash step for some proteins) and elution buffers. After that, the tags were cleaved with tobacco etch virus (TEV) protease from FL proteins using overnight incubation at +4°C. For PARP15^CAT^ R576A, D665A and D665R, TEV cleavage was not performed. For PARP15^CAT^ WT, two batches were made; one TEV-cleaved for crystallization, and one with His-tag for other experiments. Proteins were run through size exclusion chromatography (SEC) column HiLoad 16/600 Superdex 200 for PARP15^FL^ and HiLoad 16/600 Superdex S75 for PARP15^CAT^. For PARP15^FL^ additional amylose resin purification step was performed after SEC. All the collected fractions with protein, verified by running the samples on SDS-PAGE (Bio-Rad Mini-Protean TGX 4-20% gradient gel) gel, were concentrated and frozen with liquid nitrogen to small aliquots and stored at –70°C. Details of used buffers and purification of the other proteins is described in Supplementary Table 2.

### Activity assays

Activity of the proteins was evaluated using NAD^+^ conversion assay (Putt & Hergenrother, 2004; Venkannagari *et al*, 2013). For the first activity test **(Fig. 1B)** proteins were incubated with 10 mM β-NAD^+^ varying the protein concentration between 200 nM and 800 nM in 50 mM sodium phosphate (pH 7.0) buffer, quadruplicate 10 µl reactions were incubated at room temperature (RT) for 3.5 h in Revvity 384F Plus Proxiplate plate using Biosan Thermo shaker PST-60 HL plus at 400 rpm. For control reactions, we used 10 nL of 10 mM OUL232 compound that completely blocks the activity of PARP15 (Murthy *et al*, 2023). After incubation time, the remaining NAD^+^ was turned into a fluorophore by adding 2 µl of 2 M KOH and 2 µl 20% acetophenone (30% glycerol in EtOH). After 10 min, 6 µl of 100% Formic acid was added, after which the reactions were excited at 372 nm (Bandwidth 10nm) wavelength and the emission was analyzed at 444 nm wavelength (Bandwidth 20nm) with Tecan Spark plate reader. For the second activity test the reactions were incubated for 1 h at 500 nM NAD^+^ **(Fig. 3A)**.

### MARylation detection using NanoLuc-eAf1521

In the western blots, MARylation sites were revealed with macrodomain eAf1521 fused to nanoluciferase (Nluc-eAf1521), having high binding affinity for MAR groups (Boute *et al*, 2016; Nowak *et al*, 2020; Sowa *et al*, 2021). All the reactions were prepared in 10 µl volume in buffer containing 30 mM HEPES pH 7.5, 300 mM NaCl, 5% glycerol and 0.5 mM TCEP with 1 µM protein concentration and 10 µM NAD^+^. For the reactions in **Fig. 2A**, hydrolysis of MARylation was done after NAD^+^ incubation by adding 2 µM macrodomains/ADP-ribosyl hydrolases (4 µM for PARP14 macrodomain 1) to reaction followed by 1 h 15 min incubation at RT. For the first sample (PARP15^FL^, SARS-CoV-2 nsp3 MD, no NAD^+^) SARS-CoV-2 nsp3 macrodomain was added during PARP15^FL^ purification after IMAC and incubation was continued for 2 h on ice. The macrodomain was removed through amylose resin beads before continuing the protein purification with TEV cleavage.

Reactions were stopped by adding Laemmli sample buffer and boiling samples at 95°C for 5 min. Reactions were run on SDS-PAGE gels along with Precision Plus Protein all Blue standards (Bio-Rad) at 120 V. The proteins were transferred to nitrocellulose membrane using semi-dry system (Bio-Rad Trans-Blot Turbo) at 10 W for 40 min, and the complete protein transfer was confirmed by staining the gel with PageBlue™ Protein staining solution (Thermo Scientific) and the membrane with Ponceau S (Sigma) **(Fig. S7-S9)** and rinsed with (140mM NaCl, 2.7 mM KCl, 10 mM Phosphate buffer pH 7.4, 0.1 % (v/v) TWEEN®20(Sigma-Aldrich) (PBS-T, pH 7.0). Ponceau was removed from the membrane with PBS-T. The membranes were blocked using 1x TBS 1% Casein Blocker (Bio-Rad) or EveryBlot blocking buffer (Bio-Rad) followed by PBS-T wash at 4°C. 0.1 µg/ml Nluc-eAf1521 in 2.5% milk-PBS-T was incubated with the membrane before washing excess Nluc-eAf1521 away with PBS-T. The membrane was imaged with Bio-Rad’s Molecular Imager ChemiDoc^TM^ XRS+ and Image Lab^TM^ Software using Promega’s Nano-Glo® luciferase substrate in 1:1000 ratio to 10 mM sodium phosphate (pH 7.0) with Blot Hi Resolution setting.

### SEC-MALS

Light scattering detector Wyatt MiniDawn MALS and Wyatt Optilab refractive index (RI) detector were connected to a Shimadzu HPLC unit. 50 µg of each sample was run at 0.4 ml/min through Superdex 200 increase 10/300gl column with 30 mM HEPES pH 7.5, 300 mM NaCl, 5 % glycerol (v/v), and 0.5 mM TCEP in the buffer. For the analysis, Astra 7.3.2 connected to SEC-MALS unit was used.

### Mass spectrometry

Intact protein analysis **(Fig. 1C-F)** was done using a Q Exactive Plus mass spectrometer (Thermo Fisher Scientific) equipped with a heated electrospray ionization (HESI) ion source. PARP15^FL^ was incubated with NAD^+^ using 1:10 molar protein:NAD^+^ -ratio, and PARP15^CAT^ using 1:15 ratio for 2 h at room temperature. Protein samples were denatured by adding tetrafluoroacetic acid (TFA) to a final concentration of 0.1% (v/v) and injected via an Acquity UPLC system (Waters) equipped with a BioResolve RP mAb Polyphenyl Column, 450 Å, 2.7 µm, 2.1 mm x 50 mm (Waters). Approximately 1 µg of protein was used per injection. The run was set up using *Xcalibur* (Thermo) where an LC-MS method optimized for large intact proteins was selected. Data were analyzed with *BioPharma Finder* (Thermo) and deconvolved using the Respect algorithm.

For native mass spectrometry **(Fig. 5A–B, S1)**, protein samples were buffer-exchanged to 50 mM ammonium acetate using ZebaSpin (7K MWCO, 0.5 ml; Thermo Fisher Scientific) desalting columns and used in a final concentration of 10 µM. Reconstituted β-BAD sodium salt (Biolog) was added to BAD complex samples in 1–500 µM concentrations. Adjustments were done by adding 50 mM ammonium acetate. Samples were injected into the Q Exactive Plus mass spectrometer by direct infusion from a syringe with a constant flow of 10 µl/min. An MS method optimized for native proteins was used. The data were opened in and exported from *FreeStyle* (Thermo) and analyzed in *GNU Octave*. Outliers at low BAD concentrations were excluded where the calculated free concentrations were negative or occupancies were suspiciously high. The binding curves of the titration series were fitted in *Origin Pro* (Version 2018b, OriginLab Corporation) **(Fig. S6)**.

For bottom-up analyses to pinpoint the modified residues, pre-treated PARP15^FL^ WT at 20 µM was incubated with 300 µM β-NAD^+^ (15-fold molar excess) for 2 hours and then with 2 M NH_2_OH at room temperature for 1.5 hours. The bottom-up analyses were done with three different setups, the equipment used for which are detailed in Supplementary information. In general, samples were digested with a protease, separated with an HPLC system and analyzed with a mass spectrometer (Thermo Q Exactive Plus, Thermo Orbitrap Fusion Lumos Tribid or Bruker timsTOF Pro) in data-dependent tandem MS (MS/MS) mode. Automodification sites were identified in software by comparing the MS and MS/MS data against the sample sequence and allowing for a variable modification of +15.01 Da in Asp or Glu. The MS/MS spectra in which the modifications were identified are shown in **Fig. S10–S17**.

HDX-MS analyses were done using setup 3 described in Supplementary (PAL DHR Dual Head, timsTOF Pro). The robotic sampler was programmed to place 5 µl of 10 µM protein sample in 45 µl of deuterated buffer (50 mM HEPES, 150 mM NaCl, pD 7.4, in D_2_O) for four different durations (20 s, 2 min, 20 min, 2 h), after which the deuteration was quenched by adding 50 µl of quench buffer (1 M glycine-HCl, pH 2.3) and the sample was injected into the HPLC via the enzyme column. Samples with 20 s and 20 min incubation were done in triplicates. Non-deuterated samples were prepared similarly but with non-deuterated buffer (50 mM HEPES, 150 mM NaCl, pH 7.4) and no incubation time, and they were measured with both LC-MS/MS and LC-MS methods. Deuterated samples were measured with only the LC-MS method. All data were exported from *DataAnalysis* (Bruker). MS/MS data of the non-deuterated samples were compared against the sample sequences in *ProteinScape* (Bruker), whereby the signals of intact peptides were identified. Based on mass shifts of the identified peptides in LC-MS data, deuteration rates of intact peptides in deuterated samples were calculated and visualized in *DeutEx* (version 0.1.20200630). Averaged deuteration rates of each residue **(Fig 4B)** were calculated by averaging the deuteration rates of all peptides in which the residue was present.

### Macromolecular crystallography

The TEV-cleaved PARP15^CAT^ was crystallized using sitting drop vapor diffusion on an iQ plate (TTP Labtech). 75 nl of protein stock diluted to 10 mg/ml was mixed with 150 nl of crystallization solution (20% *w*/*v* PEG 3500, 0.2 M NH_4_Cl) over a reservoir containing 50 μM of the same crystallization solution using a Mosquito LCP (TTP Labtech) pipetting robot. The well was sealed with tape, and the crystallization plate was kept at room temperature. Large crystals grew in two weeks. Soaking was done by transferring a crystal to a droplet of soaking solution (20% *w*/*v* PEG 3500, 0.2 M NH_4_Cl, 50 mM HEPES, 10 mM BAD, pH 7.5) and incubating for 16 hours. Then, the crystal was transferred to cryoprotective solution (35% *w*/*v* PEG 3500, 0.2 M NH_4_Cl, 50 mM HEPES, 10 mM BAD, pH 7.5), immediately mounted in a SPINE pin and stored in liquid nitrogen.

X-ray diffraction data were collected on beamline ID30B (McCarthy *et al*, 2018) at the European Synchrotron Radiation Facility (ESRF). The images were processed using XDS (Kabsch, 2010) and cut to 1.90 Å. Molecular replacement was done with Phaser (McCoy *et al*, 2007) using pdb_00006ry4 (Korn *et al*, 2021) as the search model. Geometry restraints for BAD (CCD ID: DQV) were created with *PRODRG* (Schüttelkopf & Van Aalten, 2004) using idealized coordinates downloaded from the PDB as the input file. The structure was refined with *Refmac*5 (Murshudov *et al*, 2011) run via the *CCP*4*i*2 interface (Potterton *et al*, 2018; Winn *et al*, 2011). Riding hydrogens were used in refinement and omitted from the output file. Edits to the model between refinement rounds were done with *Coot* (Emsley *et al*, 2010). The finalized structure was deposited in the Protein Data Bank (PDB) under the accession code pdb_00009tcb.

### Immunoprecipitation assays

To confirm dimerization HEK293 cells were transfected with plasmids encoding either HA-PARP15.1-HY or HA-PARP15.1-HY-DR and co-transfected with plasmids encoding GFP or GFP-PARP15.1-HY via the calcium phosphate precipitation method. After 24 h cells were lysed using TAP-lysis buffer (50 mM Tris, pH 7.5; 150 mM NaCl; 1 mM EDTA; 10% Glycerol; 1% NP-40; 2 mM TCEP; PIC) and lysates cleared by centrifugation at 4°C for 30 min. GFP or the GFP-PARP15.1-HY fusions were enriched by incubation with 5 µl GFP-Trap magnetic agarose beads at 4°C for 1 h. Afterwards the beads were washed 4 times with TAP-lysis buffer (increased NaCl concentration of 250 mM). Immunoprecipitated material as well as whole cell lysates were separated by SDS-PAGE and analyzed via immunoblotting using specific antibodies.

To evaluate the MARylation status of PARP15 variants, stable U2OS-Flp-In TRex cells were induced with 0.5 µl/ml doxycycline for the expression of either GFP-PARP15 WT, GFP-PARP15-H559Y or GFP-PARP15-D665R overnight. The next day, cells were harvested using RIPA lysis buffer (10 mM Tris, pH 7.4; 150 mM NaCl; 1% NP-40; 1% DOC; 0.1% SDS; PIC) and GFP-fusion proteins enriched as described before. After four wash steps with RIPA lysis buffer, whole cell lysates and immunoprecipitated material were separated by SDS-PAGE and immunoblot analysis was performed using specific antibodies. Antibodies used in these studies: anti-HA (BioLegend, clone 16B12), anti-GAPDH (sc-32233), anti-GFP (Proteintech, 66002-1-lg), anti-PARP15 antibodies (Proteintech, 18126-1-AP), anti-Poly/Mono-ADP-ribose (E6F6A) antibody (Cell Signaling, 83732S), goat-anti-rabbit-HRP (Jackson Immunoresearch, 111-035-144), goat-anti-mouse-HRP (Jackson Immunoresearch, 115-036-068).

### Prediction of protein structures and graphical design

AlphaFold3 server was used for predicting the protein structures (Abramson *et al*, 2024). Graphs for NAD^+^ hydrolysis assay and SEC-MALS were created with GraphPad Prism version 10.1.2. for Windows, GraphPad Software, Boston, Massachusetts USA, www.graphpad.com. PyMOL was used for presentation of the crystal structure and AlphaFold predictions (Schrödinger, L., & DeLano, W. (2020). *PyMOL*. Retrieved from http://www.pymol.org/pymol).

## Acknowledgements

The use of the facilities and expertise of the Biocenter Oulu Structural Biology core facility (a member of Biocenter Finland, Instruct-ERIC Centre Finland and FINStruct), Proteomics and Protein Analysis core facility (a member of Biocenter Finland) and Biocenter Oulu sequencing center are gratefully acknowledged.

We acknowledge the European Synchrotron Radiation Facility for provision of synchrotron radiation facilities and we would like to thank the staff of the ESRF and EMBL Grenoble for assistance and support in using beamline ID30B under the proposal number MX2553.

This work benefited from access to BIOCEV, an Instruct-ERIC Centre. Financial support was provided by Instruct-ERIC (internship APPID 3678 and PID 33245). In particular, we thank Dr. Petr Pompach from BIOCEV for his guidance in the structural mass spectrometry facility.

We thank Men Thi Hoai Duong, Juho Alaviuhkola and Naveed Ahmad for help with protein production. Expression construct PARP15 macrodomain 2 construct was a generous gift from Structural Genomics Consortium (Stockholm, Sweden).

## CRediT authorship contribution statement

**Anna Tuovinen:** Investigation, Formal analysis, Visualization, Writing-original draft; **Johan Pääkkönen:** Methodology, Investigation, Data curation, Formal analysis, Visualization, Writing-review and editing; **Mirko Maksimainen:** Conceptualization, Supervision; **Lea Hirschen:** Investigation; **Heli I. Hentilä**: Investigation; **Marie Tauscher:** Investigation; **Bernhard Lüscher**: Methodology, Supervision, Writing-review and editing; **Carlos Vela-Rodríguez**: Methodology, Writing-review and editing, Supervision; **Patricia Korn**: Writing-review and editing, Supervision; **Lari Lehtiö:** Conceptualization, Validation, Writing-review and editing, Supervision.

## Data availability

Atomic coordinates and structure factors will be available at the Protein Data Bank with the ID pdb_00009tcb. Raw diffraction data will be available at fairdata.fi (https://doi.org/10.23729/fd-ea8ba18c-8148-35cf-af5b-033f9d8f056a). Other study data are included in the article and supporting information.

